# Common and unique effects of HD-tDCS to the social brain across cultural groups

**DOI:** 10.1101/408799

**Authors:** A. K. Martin, P. Su, M. Meinzer

**Affiliations:** The University of Queensland, UQ Centre for Clinical Research, Brisbane, Australia; Durham University, Department of Psychology, Durham, UK; University of Greifswald, Department of Neurology, Greifswald, Germany

**Keywords:** self-construal, visual perspective taking, self-reference effect, mPFC, rTPJ, cross-cultural

## Abstract

**ABSTRACT:** Cultural background influences social cognition, however no study has examined brain stimulation differences attributable to cultural background. 104 young adults [52 South-East Asian Singaporeans (SEA); 52 Caucasian Australians (CA)] received anodal high-definition transcranial direct current stimulation (HD-tDCS) to the dorsomedial prefrontal cortex (dmPFC) or the right temporoparietal junction (rTPJ). Participants completed tasks with varying demands on self-other processing including visual perspective taking and episodic memory with self and other encoding. At baseline, SEA showed greater self-other integration than CA in the level one (line-of-sight) VPT task as indexed by greater interference from the alternate perspective. Anodal HD-tDCS to the dmPFC resulted in the CA performing closer to the SEA during egocentric perspective judgements. Baseline performance on level two (embodied rotation) VPT task and the self-reference effect in memory (SRE) was comparable between the two groups. In the combined sample, HD-tDCS to the rTPJ decreased the interference from the egocentric perspective during level two VPT and dmPFC HD-tDCS removed the SRE in episodic memory. Stimulation effects were comparable when baseline performance was comparable. When baseline performance differed, stimulation differences were identified. Therefore, social cognitive differences due to cultural background are an important consideration in social brain stimulation studies.

**HIGHLIGHTS:**

- Compared with Caucasians, South-East Asians were influenced by the alternate perspective to a greater extent during level one visual perspective taking
- Anodal HD-tDCS to the dmPFC shifted Caucasians closer to the baseline performance of South-East Asians
- Anodal HD-tDCS to the dmPFC removed the self-reference effect in episodic memory in both cultural groups
- Anodal HD-tDCS to the dmPFC reduced overall memory performance in the South-East Asians but not in the Caucasian group
- Anodal HD-tDCS to the rTPJ reduced egocentric interference in a level two visual perspective taking task in both cultural groups

## 1. INTRODUCTION

Social cognition refers to the broad range of cognitive processes that facilitate social behaviour. These cognitive processes are shaped by societal influences such as the cultural background present throughout development. For example, how the notion of self is defined in relation to others is thought to be culturally specific. People from East Asia, who are generally organized in more prototypically collectivistic societies (Markus & Kitayama, 1991), may process social information in a more integrative fashion, whereby the self is defined in relation to others rather than a distinct construct. It has also been suggested that the East Asian sense of self is more relational, whereby the demands of the situation override internalized attributes (Heine, 2001). On the contrary, people from more individualistic oriented cultural backgrounds, such as Western societies, tend to position the self as distinct from others (Markus & Kitayama, 1991). Despite studies demonstrating differences in social cognition and the underlying social brain dependent on cultural background (Han et al., 2013), no study to date has investigated whether non-invasive brain stimulation is influenced by cultural background.

Key components of social cognition are the ability to both integrate and distinguish between processes, or representations, relevant to the self or another person. Such ability is considered key for higher-order social cognitive functioning such as empathy and theory of mind (ToM). Individualistic or collectivistic self-construal has been shown to influence social cognition, in particular those that manipulate self-other processing (Han et al., 2013; Kessler, Cao, O’Shea, & Wang, 2014; Kühnen, 2009; Sparks, Cunningham, & Kritikos, 2016; Wu & Keysar, 2007). Visual perspective taking is considered a lower-level cognitive process thought to be relevant for empathy and ToM with measurable demands on self-other integration and distinction (Apperly & Butterfill, 2009). Cultural background influences VPT performance. For example, Wu & Keyser (2007) found that Chinese students were better than American Caucasian students at taking their partner’s perspective in a communication task. Using a VPT task requiring line of sight judgements (VPT level one), Kessler & colleagues (2014) demonstrated a greater egocentric bias in those from a Western culture compared to those from an East Asian cultural background with no difference on a VPT task requiring mental rotation (VPT level two).

Self-other processing is relevant across several cognitive domains including episodic memory. The self-reference effect (SRE) refers to the phenomenon whereby people recall or recognise information with greater accuracy if encoded in relation to the self (Symons & Johnson, 1997). Again, this phenomenon is culturally specific, with those from an East Asian background showing less of a self-reference effect when the other is a close family member. However, the SRE is comparable to that of Western participants when the other is not a personal acquaintance (Sparks et al., 2016). We would therefore expect similar brain-behaviour associations for self and other encoded words in CA and SEA people, if the other is a familiar but non-family member. Consequently, excitatory stimulation of the social brain should result in comparable modulation of the self-reference effect in both groups.

Two key hubs of the social brain associated with the integration and distinction between self and other are the dorsomedial prefrontal cortex (dmPFC) and the right temporoparietal junction (rTPJ) (Ferrari et al., 2016; Payne & Tsakiris, 2017; Santiesteban, Banissy, Catmur, & Bird, 2012; Wang, Callaghan, Gooding-Williams, McAllister, & Kessler, 2016; Wittmann et al., 2016). Recently, we have demonstrated that high-definition transcranial direct current stimulation (HD-tDCS) to the dmPFC and rTPJ, can produce regional and task specific effects on self-other processes (Martin, Dzafic, Ramdave, & Meinzer, 2017; Martin, Huang, Hunold, & Meinzer, 2018). Excitatory stimulation to the dmPFC increased the integration of the other (allocentric) perspective into the self (egocentric) perspective across explicit VPT tasks and removed the SRE in episodic memory. Excitatory stimulation to the rTPJ resulted in a selective effect on reducing the egocentric interference when making an embodied rotation into an alternate perspective (Martin, Huang, et al., 2018).

Cultural background has also been demonstrated to shape the underlying brain-behaviour relationship, especially in the medial prefrontal cortex (mPFC; Han & Ma, 2014; Han et al., 2013; Harada, Li, & Chiao, 2010; Ma, Bang, et al., 2014; Ma, Wang, et al., 2014; Paige, Ksander, Johndro, & Gutchess, 2017; Park & Huang, 2010; Ray et al., 2010; Zhu, Zhang, Fan, & Han, 2007). Therefore, cultural background may be a potential source of variability in the effects of non-invasive brain stimulation, specifically stimulation of the mPFC. The aims of the present study are to replicate previous effects of HD-tDCS on social cognition (Martin, Huang, et al., 2018) and identify unique differences due to cultural background. As culture is not a homogenous construct, self-construal will be assessed in the South-East Asian (SEA) participants using a multi-dimensional measure (Vignoles et al., 2016) and exploratory analyses conducted to further assess culturally specific effects of brain stimulation. Self-construal has previously been shown to influence social cognition (Colzato, de Bruijn, & Hommel, 2012; Cross, Morris, & Gore, 2002; Kühnen, 2009) and underlying brain-behaviour relationships, especially the mPFC (Chen, Wagner, Kelley, & Heatherton, 2015; Li et al., 2018; Liu, Wu, Petti, Wu, & Han, 2018; Ma, Wang, et al., 2014; Ray et al., 2010). However, no study to date has investigated the relationship between self-construal and focal brain stimulation response.

In line with previous literature (Kessler et al., 2014; Sparks et al., 2016), we hypothesise that compared with Caucasian Australians (CA), SEA will show a greater congruency effect in the level one VPT tasks but similar baseline performance on level two VPT and similar extent of the SRE in episodic memory. We hypothesise that anodal stimulation to the dmPFC and the rTPJ will have a comparable effect in SEA as those identified previously in CAs (Martin, Dzafic, et al., 2017; Martin, Huang, et al., 2018). Specifically, dmPFC stimulation will result in greater congruency effects across both level one and two tasks and a reduction/removal of the SRE. Anodal stimulation to the rTPJ will result in reduced interference from the egocentric perspective during the level two VPT task only.

## 2. METHOD

### 2.1 Participants

Fifty-two participants from Singapore, residing in Brisbane, Australia were recruited and stratified to receive either dmPFC or rTPJ anodal HD-tDCS in a sham-controlled, double-blinded, repeated-measures design. The groups were matched for age and gender. A cohort of 52 Caucasian Australians (CA) served as a comparison group (data already published see Martin, Huang, et al., 2018). The CA and South-East Asian (SEA) groups were matched for age (23.4 v 23.2yrs, BF_10_= 0.22) and gender (13 M/F for each group and stimulation site). A power analysis was computed using G*Power 3.1 indicating a sample size of 96 as adequate power (0.8) with alpha set at 0.05. A sample of 104 was used to guard against dropout and maintain identical sample sizes with a previous study in Caucasian Australians only (Martin, Huang, et al., 2018). All participants were tDCS-naïve and had no history of psychiatric or neurological disorder and reported no psychoactive substance or medication use. All participants provided written consent in accordance with the Declaration of Helsinki (1991; p1194), they completed a safety screening form to ensure they were safe to participate. All were provided with a small monetary compensation. The ethics committee of The University of Queensland granted ethical approval.

### 2.2 Baseline Testing

Participants completed the Hospital Anxiety and Depression Scale (HADS) (Zigmond & Snaith, 1983) and the Autism-Spectrum Quotient (Baron-Cohen, Wheelwright, Skinner, Martin, & Clubley, 2001). Baseline cognitive testing was performed to ensure participants were within age appropriate norms. This included the Stroop Test, phonemic and semantic verbal fluency, and the following tests from the CogState^®^ computerized test battery (http://cogstate.com): International shopping list, identification test, one-back, two-back, set-switching task, continuous paired-associates learning test, social-emotional cognition test, and the International shopping list – delayed recall.

### 2.3 Transcranial Direct Current Stimulation (tDCS)

Stimulation was administered using the one-channel direct current stimulator (DC-Stimulator Plus^®^, NeuroConn) with two concentric rubber electrodes (Bortoletto, Rodella, Salvador, Miranda, & Miniussi, 2016; Gbadeyan, Steinhauser, McMahon, & Meinzer, 2016). The centre electrode was 2.5cm in diameter for both the dmPFC and rTPJ sites. The return electrode was 9.2/11.5 cm inner/outer in diameter for the dmPFC site, whereas a smaller electrode of 7.5/9.8 cm was used for the rTPJ site, due to the position of the right ear. Current modelling and safety information for these montages are presented in detail in previous studies (Gbadeyan et al., 2016; Martin, Huang, Hunold, & Meinzer, 2017; Martin, Huang, et al., 2018). Electrodes were attached using an adhesive conductive gel (Weaver Ten20^®^ conductive paste) and held firmly in place using an elastic EEG cap. Both dmPFC and rTPJ sites were identified using the 10-20 International EEG system. The dmPFC was located by finding 15% of the distance from the Fz towards the Fpz. The rTPJ was located by finding CP6. For both sham and anodal stimulation conditions, the current was ramped up to 1mA over 40 secs before ramping down over 5 secs. In the anodal stimulation condition the current remained at 1mA for 20 minutes prior to ramping down. Researchers used the “study-mode” of the DC-Stimulator to achieve double-blinding. Stimulation sessions were at least 3 days apart to avoid carry-over effects. Stimulation order was counterbalanced in both CA and SEA groups so that half received active stimulation in the first session and half received active stimulation in the second session.

### 2.4 Visual Perspective Taking Task

The visual perspective taking (VPT) task has been described in detail previously (Martin, Dzafic, et al., 2017; Martin, Huang, et al., 2018; Martin et al., 2019). In brief, three separate tests were administered measuring implicit, explicit level one and explicit level two. All levels involved a street scene with either an avatar or traffic light with 1-4 tennis balls present. The avatar/traffic light was in 1 of 3 locations on the street; far, middle, or near (see Martin, Huang, et al., 2018 for more details). The traffic light was a directional control intended to direct attention in a similar manner to the avatar but crucially without the ability to hold a perspective. This was of particular interest for inducing an implicit VPT effect, where participants should respond slower when the scene is incongruent with the hypothetical view of the avatar but not the traffic light (Apperly & Butterfill, 2009; Samson, Apperly, Braithwaite, Andrews, & Bodley Scott, 2010). There were 176 trials in each VPT level. For the implicit task this was divided into 4 equal groups of 44 across the four conditions (avatar congruent, avatar incongruent, light congruent, light incongruent). For the two explicit tasks, the trials were divided into 8 equal groups of 22 to accommodate both egocentric and allocentric conditions. All conditions were balanced for number and location of tennis balls, and location of avatar and traffic light. The trials were randomised into 4 versions to avoid order effects and provide different versions between sessions. Participants always completed the VPT tasks in the following order; implicit, explicit level one, and explicit level two. Participants were instructed to answer as quickly and accurately as possible. A fixation cross was presented for 500msec prior to each trial. In the explicit tasks, a screen with You or Other was presented for 750msecs after the fixation cross to signal whether the perspective to take was the egocentric or allocentric.

The task was designed to keep errors low and therefore only response times were of interest in the current study. The main outcome of interest was the congruency effect (incongruent – congruent) for both egocentric and allocentric conditions. The avatar and traffic light factor was only of interest in the implicit VPT task and was collapsed across for the explicit tasks, similar to previous studies (Martin, Dzafic, et al., 2017; Martin, Huang, et al., 2018; Martin et al., 2019; Santiesteban, Catmur, Hopkins, Bird, & Heyes, 2014).

### 2.5 Implicit VPT

Initially, participants were asked to answer as quickly and as accurately as possible “How many tennis balls can you see?” The answer was between 1-4. This was considered an implicit task as participants only responded from the egocentric perspective and were not directed to the perspective of the avatar in the scene.

### 2.6 Explicit level one

In the level one task, participants were instructed to answer how many tennis balls they could see or how many the avatar could see, or if a traffic light in the scene, how many balls the light would shine on directly? The answer was between 1-4 for the congruent and incongruent egocentric conditions and congruent allocentric conditions. To maintain four options, the incongruent allocentric condition contained between 0-3 tennis balls.

### 2.7 Explicit level two

In the level two task, participants were now required to make a laterality judgement and answer “can you see more tennis balls on the left/right/same?” When taking the allocentric perspective, participants answered whether the avatar could see more balls on the left/right/same or whether the traffic light would directly shine light on more balls on the left/right/same?

### 2.8 Self-Referential Memory Task

Prior to the VPT, participants completed a modified version of the Reading the Mind in the Eyes Test (RMET) (Baron-Cohen, Wheelwright, Hill, Raste, & Plumb, 2001), data published elsewhere (Martin, Huang, et al., 2017; Martin, Su, & Meinzer, 2018). The RMET requires participants to select one word from four multiple-choice options to describe the emotion/ mental state that best describes the expression from the eye region of the face. The control condition required selecting the age and sex of the eyes. Following this they were asked how often they thought or felt that way (self-encoded) or how often they thought Barack Obama would feel or think that way (other-encoded). To ensure participants were familiar with Barack Obama, participants viewed a 5-minute documentary on his life prior to testing. To encourage engagement in the task, participants were told that their scores would be compared against those who had worked alongside Barack Obama.

Following the completion of the VPT tasks, participants performed a recognition memory task for the mental attribution words from the RMET. The options included 38 correct RMET words, 38 incorrect distractor words from RMET and 38 novel unseen words. Participants were presented with the word and asked if they had seen it in the eyes. They could respond in a confident manner (definitely did/ did not) or in a less confident manner (probably did/ did not). Scoring was 2 for correct confident response through to −2 for an incorrect confident response.

### 2.9 Source Memory Task

If participants identified that they had seen the mental attribution in the eyes, they were then asked “was it on a male or female face?” They could respond in a confident manner (definitely male/female) or less confident (probably male/female). Scoring was identical to the SRE task above.

A schematic description of all tasks and stimulation procedures is presented in a previous study (Martin, Dzafic, et al., 2017).

### 2.10 Self-Construal Style

The SEA participants also completed a self-construal scale (Vignoles et al., 2016) to measure self-reported interdependence or independence as to how the self is perceived in relation to others. The 22-item self-construal scale can be decomposed into 7 dimensions; Self-reliance vs Dependence on others (SRvDO), Connection to others vs Self-containment (COvSC), Difference vs Similarity (DvS), Self-interest vs Commitment to others (SIvCO), Consistency vs Variability (CvV), Receptiveness to influence vs Self-direction (RIvSD), Self-expression vs Harmony (SEvH). Participants responded by indicating their agreement or disagreement on a 9-point scale ranging from 1 (not at all) to 9 (exactly). Scores were reversed where necessary so that higher scores reflected greater independent and lower scores greater interdependent self-construal style. Self-construal scores were then correlated with stimulation effects across all self-other tasks. This was predominantly of interest for the SRE in episodic memory, in line with previous research showing self-construal was associated with differences in underlying social brain regions during self-referential tasks (Chen et al., 2015; Chiao et al., 2009).

### 2.11 Adverse Effects and Blinding

Following each session participants were asked if they experienced any adverse effects (Alam, Truong, Khadka, & Bikson, 2016) and their mood was assessed prior to and after each stimulation session using the Visual Analogue of Mood Scales (VAMS) (Folstein & Luria, 1973). Participants also stated which session they thought was active stimulation to assess whether blinding was effectively achieved.

### 2.12 Current Modelling

Current modelling was conducted for both the dmPFC and rTPJ stimulation sites and is presented in previous manuscripts (Martin, Huang, et al., 2017, 2018). Briefly, peak current field strengths (0.36 V/m) were identified at the dmPFC [MNI: 0 54 33] and rTPJ (0.59 V/m) [MNI: 60 −54 13]. Focal current delivery to the dmPFC and rTPJ was demonstrated with physiologically effective current strengths and also compares favourably to previous current modelling studies (see Martin, Huang, et al., 2018 for further details).

### 2.13 Statistical Analyses

All analyses were completed using Bayesian statistics in JASP 0.8.6. We used the Bayes Factor (BF) to quantify evidence for the null or an alternate model. The BF_10_ refers to the likelihood of the data for a particular model. For example, a BF_10_ = 8 equates to data that is 8 times more likely given the alternate model than the null model. Bayes Factors should be interpreted in a linear fashion, with larger BF_10_ providing greater evidence for the alternate mode, but some thresholds have been proposed for ease of interpretation. BF_10_ 1-3 is support for the alternate model but should be considered preliminary; BF_10_ 3-10 should be considered moderate evidence, and any BF_10_ > 10 considered strong evidence. The inverse is true for the null model with BF_10_ 0.3-1 providing preliminary evidence, BF_10_ 0.1-0.3 moderate, and BF_10_ < 0.1 strong evidence. Although not a consistent match in all cases, preliminary evidence usually translates to frequentist p-values between 0.01-0.05, moderate evidence 0.005-0.01, and strong evidence to <0.005 (Benjamin et al., 2018). Default priors were used across all analyses as recommended (Wagenmakers et al., 2018).

Repeated measures analysis of variance (RM-ANOVA) were computed for congruency effect with stimulation type and perspective as within participant factors and cultural group and stimulation site as between participant factors.

## 3. RESULTS

Performance on all tasks are presented in Table 1.

**Table 1.** Performance on the Visual Perspective Taking and Episodic and Source Memory tasks across stimulation site, cultural group, and stimulation type. Response times refer to the difference between incongruent and congruent trials in milliseconds (msecs).

|  | dmPFC |  |  |  | rTPJ |  |  |  |
| --- | --- | --- | --- | --- | --- | --- | --- | --- |
|  | South-East Asians |  | Caucasian<br>Australians |  | South-East Asians |  | Caucasian<br>Australians |  |
|  | Sham<br>Mean<br>(sd) | Anodal<br>Mean<br>(sd) | Sham<br>Mean<br>(sd) | Anodal<br>Mean<br>(sd) | Sham<br>Mean<br>(sd) | Anodal<br>Mean<br>(sd) | Sham<br>Mean<br>(sd) | Anodal<br>Mean<br>(sd) |
| *Level two VPT* |  |  |  |  |  |  |  |  |
| Ego CE RT (msecs) | 167.34<br>(179.30) | 157.91<br>(145.57) | 120.58<br>(139.41) | 195.88<br>(134.74) | 106.12<br>(208.50) | 195.32<br>(200.32) | 138.76<br>(147.85) | 131.12<br>(190.67) |
| Allo CE RT (msecs) | 233.27<br>(225.08) | 245.77<br>(173.26) | 249.81<br>(141.56) | 223.16<br>(163.67) | 315.25<br>(269.75) | 213.26<br>(237.70) | 244.31<br>(157.87) | 108.25<br>(167.26) |
| *Level one VPT* |  |  |  |  |  |  |  |  |
| Ego CE RT (msecs) | 159.75<br>(111.70) | 186.56<br>(163.94) | 74.31<br>(79.09) | 122.75<br>(116.05) | 138.91<br>(163.55) | 176.65<br>(129.04) | 153.72<br>(136.35) | 130.87<br>(128.65) |
| Allo CE RT (msecs) | 185.24<br>(157.54) | 236.76<br>(142.75) | 184.35<br>(100.88) | 149.38<br>(145.32) | 275.76<br>(151.11) | 218.18<br>(199.06) | 208.32<br>(106.50) | 184.13<br>(119.40) |
| *Implicit VPT* |  |  |  |  |  |  |  |  |
| Avatar CE RT (msecs) | 14.90<br>(22.23) | 14.47<br>(18.92) | 16.29<br>(18.98) | 12.48<br>(24.33) | 17.64<br>(30.69) | 11.03<br>(24.45) | 8.03<br>(22.72) | 9.97<br>(25.05) |
| Light CE RT (msecs) | 1.64<br>(22.86) | -3.90<br>(26.86) | -4.70<br>(20.30) | -6.74<br>(19.85) | 0.10<br>(32.60) | -1.84<br>(23.49) | -0.09<br>(17.82) | -6.05<br>(20.43) |
| *Episodic Memory* |  |  |  |  |  |  |  |  |
| Self-Encoded | 1.03<br>(0.42) | 0.50<br>(0.90) | 0.89<br>(0.53) | 0.76<br>(0.58) | 0.44<br>(0.62) | 0.56<br>(0.69) | 0.64<br>(0.67) | 0.65<br>(0.66) |
| Other-Encoded | 0.66<br>(0.60) | 0.58<br>(0.61) | 0.58<br>(0.68) | 0.72<br>(0.59) | 0.47<br>(0.66) | 0.62<br>(0.63) | 0.64<br>(0.71) | 0.50<br>(0.72) |
| *Source Memory* |  |  |  |  |  |  |  |  |
| Self-Encoded | 0.62<br>(0.42) | 0.50<br>(0.50) | 0.60<br>(0.33) | 0.64<br>(0.43) | 0.51<br>(0.54) | 0.50<br>(0.51) | 0.61<br>(0.46) | 0.54<br>(0.43) |
| Other-Encoded | 0.48<br>(0.49) | 0.51<br>(0.45) | 0.68<br>(0.48) | 0.66<br>(0.35) | 0.46<br>(0.53) | 0.69<br>(0.48) | 0.57<br>(0.44) | 0.68<br>(0.44) |

VPT= Visual Perspective Taking Task; dmPFC= dorsomedial prefrontal cortex; rTPJ= right temporoparietal junction; sd= standard deviation; CE= congruency effect; msecs= milliseconds; Ego= Egocentric; Allo= Allocentric.

### 3.1 Visual Perspective Taking (VPT)

#### 3.1.1 Level one VPT

A main effect of Perspective was identified, BF_10_ = 42751.473, η_p_^2^ = 0.20 such that the congruency effect was greater when taking the allocentric perspective compared to the egocentric perspective. This was comparable across the Cultural Groups, BF_10_ = 0.149, η_p_^2^ = 0.00, although an overall difference in congruency effect (egocentric & allocentric combined) between the two cultural groups was identified, BF_10_ = 2.436, η_p_^2^ = 0.06 (see Figure 1A). Evidence for an interaction between Cultural Group x Stimulation site x Stimulation Type x Perspective was identified, BF_10_ = 2.221, η_p_^2^ = 0.06. As we previously identified site-specific effects on level 1 VPT, we next computed separate 3-way ANOVAs for each stimulation site.

**Figure 1.**
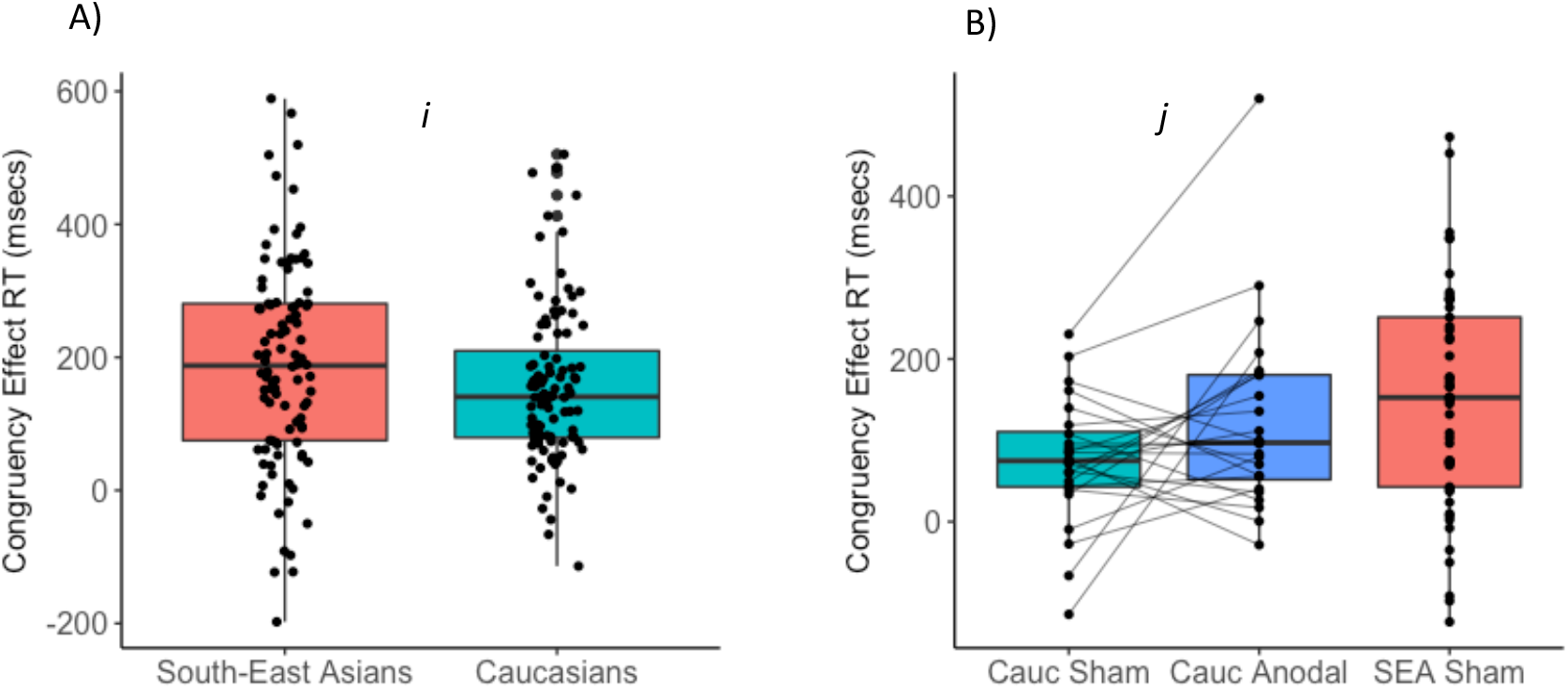
A) Level one VPT (pooled across both sham and anodal sessions). South-East Asian (SEA) participants had a greater congruency effect than Caucasian Australians (CA) demonstrating that they were more influenced by the perspective other than the one currently required for task demands (*i)*, BF_10_ = 2.436. B) During the egocentric condition of the VPT level one task, SEA showed a greater interference effect due to the allocentric perspective and in the CA group, anodal stimulation to the dmPFC shifted performance closer to the SEA group (*j*), BF_10_ = 1.012, such that the allocentric perspective influenced egocentric performance to a greater extent. The boxplot displays the median and the interquartile range (IQR). The whiskers extend to the most extreme datapoint within ±1.5*IQR.

In the rTPJ stimulation group, there was no support for a three-way interaction between Cultural Group x Stimulation Type x Perspective, BF_10_ = 0.659, η_p_^2^ = 0.05. No support was found for the two-way interactions between Stimulation x Cultural Group, BF_10_ = 0.222, η_p_^2^ = 0.003 and for Stimulation Type x Perspective, BF_10_ = 0.429, η_p_^2^ = 0.05. There was also no support for the main effect of stimulation, BF_10_ = 0.236, η_p_^2^ = 0.02.

In the dmPFC stimulation group, support for a three-way interaction between Cultural Group x Stimulation x Perspective was identified, BF_10_ = 1.041, η_p_^2^ = 0.07. Separate 2-way ANOVAs were computed for each cultural group. In the CA group, we identified preliminary evidence for a site-specific effect of dmPFC stimulation (stim site x stim type interaction BF_10_ = 1.723) and for anodal stimulation to the dmPFC increasing the congruency effect during egocentric condition only, BF_10_ = 1.012, δ = 0.45 (see Martin, Huang, et al., 2018 for further details). In the SEA group there was no Stimulation x Perspective interaction, BF_10_ = 0.309, η_p_^2^ = 0.01, nor was there an overall effect of stimulation, BF_10_ = 0.779, η_p_^2^ = 0.12.

Therefore, the evidence supported a null effect for anodal stimulation to either the dmPFC or the rTPJ in the SEA group. Whereas, in the CA group, anodal stimulation to the dmPFC increased the interference from the allocentric perspective during egocentric perspective taking, shifting the performance of the CA closer to that of the SEA group (see Figure 1B).

#### 3.1.2 Level two VPT

There was a four-way interaction between Stim Type x Stimulation Site x Perspective x Cultural Group was identified, BF_10_ = 2.025, η_p_^2^ = 0.02. As we previously identified a specific effect for rTPJ stimulation on the allocentric congruency effect (see Martin, Huang, Hunold, & Meinzer, 2018), we next calculated separate 3-way ANOVAs for both the rTPJ and dmPFC stimulation sites.

In the rTPJ stimulation group, a significant Stim Type x Perspective was identified, BF_10_ = 12.428, η_p_^2^ = 0.15 and cultural group had no effect, Stim Type x Perspective x Cultural Group, BF_10_ = 0.288, η_p_^2^ = 0.01. Paired t-tests showed a significant reduction of allocentric congruency effect after rTPJ stimulation (280.50 v 161.83 msecs, BF_10_ = 17.850, δ = 0.48). Stimulation had no effect on the egocentric congruency effect (122.11 v 163.87 msecs), BF_10_ = 0.318, δ = −0.18).

After dmPFC stimulation, a significant Stim Type x Perspective x Cultural Group interaction was identified, BF_10_ = 1.046, η_p_^2^ = 0.06. Therefore, we performed two separate 2-way ANOVAs for SEA and CAs. The results for the CAs have previously been reported (Martin, Huang, Hunold, & Meinzer, 2018). Briefly, anodal stimulation to the dmPFC increased the interference from the allocentric perspective during egocentric perspective judgements as indexed by an increased egocentric congruency effect. In the SEA group, there was no Stim x Perspective interaction, BF_10_ = 0.302, η_p_^2^ = 0.004, nor a Stimulation effect overall, BF_10_ = 0.216, η_p_^2^ = 0.00. However, a main effect of perspective was identified, BF_10_ = 2.040, η_p_^2^ = 0.15, such that the allocentric congruency effect was greater (239.52 msecs) than the egocentric congruency effect (162.62 msecs) in the SEA group. Therefore, anodal stimulation to the dmPFC resulted in a shift of performance in the CA group closer to that of the SEA at baseline.

**Figure 2.**
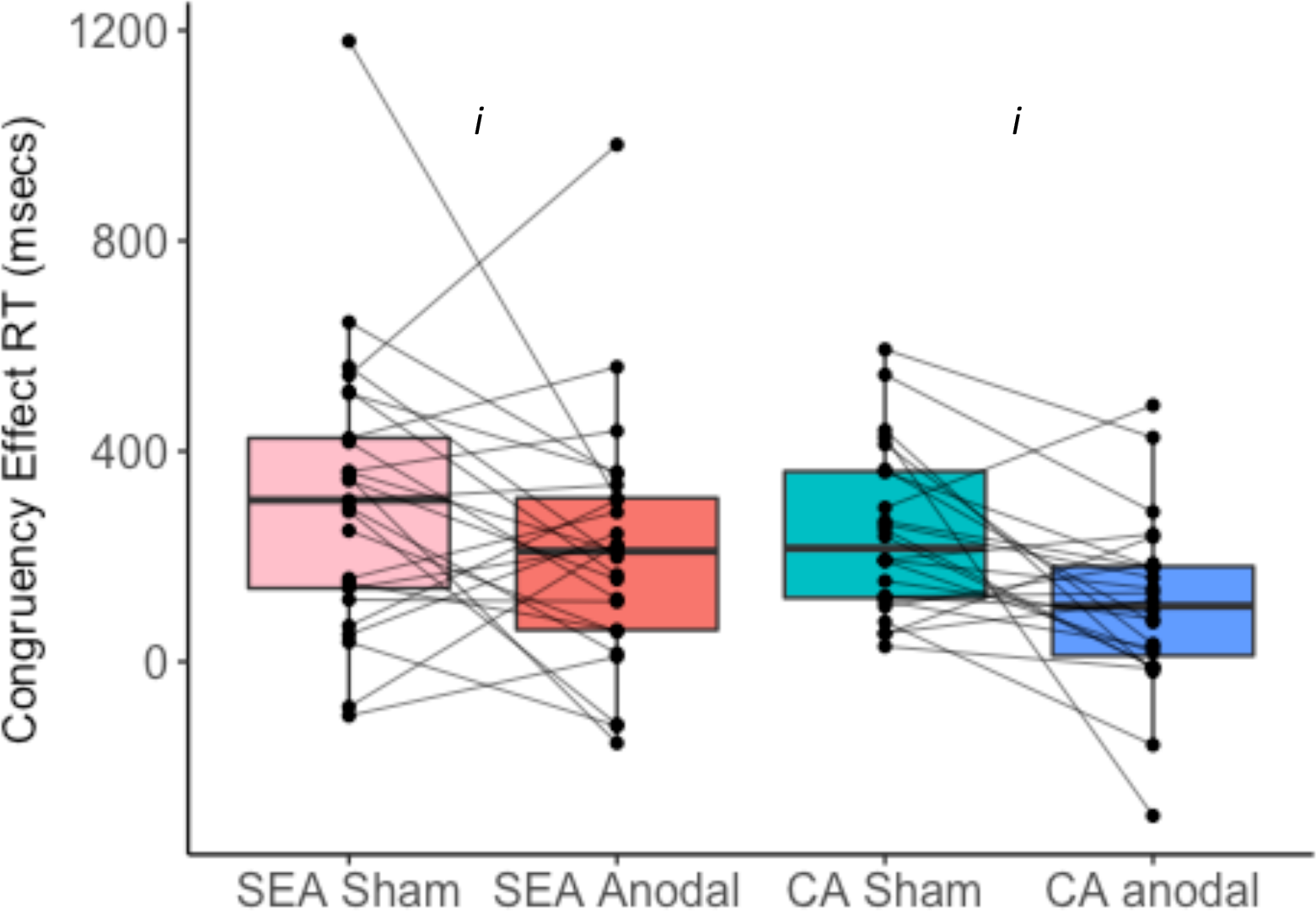
Level two VPT. Anodal stimulation to the right TPJ reduced the congruency effect due to interference from the egocentric perspective during the allocentric condition of the level two VPT (*i*), BF_10_ = 12.428. This was consistent across both cultural groups, BF_10_ = 0.288. The boxplot displays the median and the IQR. The whiskers extend to the most extreme datapoint within ±1.5*IQR. CA= Caucasian Australian; SEA= South-East Asian

#### 3.1.3 Implicit VPT

A main effect of Agent was identified, BF_10_ = 2.643e+9, ηp^2^ = 0.37. Simple effects analysis identified a significant difference between congruent and incongruent scenes when an avatar was in the scene, BF_10_ = 1.299e+10, δ = −1.12, but not when a traffic light was in the scene, BF_10_ = 0.467, δ = 0.23. There was no difference in the implicit VPT effect between the two cultural groups, BF_10_ = 0.143, η_p_^2^ = 0.01. Stimulation had no overall effect on implicit VPT, BF_10_ = 0.162, η_p_^2^ = 0.001, nor a site-specific effect, BF_10_ = 0.209, η_p_^2^ = 0.00. Cultural group had no effect on stimulation response, BF_10_ =0.154, η_p_^2^ = 0.001 nor on the site-specific effect, BF_10_ = 0.245, η_p_^2^ = 0.001.

### 3.2 Self-Reference Effect on Episodic Memory

A SRE was identified for episodic memory, BF_10_ = 6.721, η_p_^2^ = 0.08 whereby self-encoded words were better remembered than other-encoded words. The SRE was comparable between the two cultural groups, BF_10_ = 0.214, η_p_^2^ = 0.000 demonstrating comparable baseline performance. The site-specific effects of stimulation were different between the two cultural groups, as indexed by support for the Stimulation Type x Stimulation Site x Cultural Group interaction, BF_10_ = 10.204, η_p_^2^ = 0.06. In addition, support was found for a Stimulation Type x Stimulation Site x Agent interaction, BF_10_ = 2.526, η_p_^2^ = 0.08 that did not differ between the cultural groups, BF_10_ = 0.27, η_p_^2^ = 0.00.

First, as an interaction was identified for a site-specific and agent-specific effect of stimulation that was comparable across cultural groups, each stimulation site was analyzed separately collapsed across cultural group. For the dmPFC stimulation group, a Stimulation Type x Agent interaction was identified, BF_10_ = 6.542, η_p_^2^ = 0.21. Simple effects analyses identified a SRE during sham, BF_10_ = 746.63, δ = 0.85 that was removed after anodal dmPFC stimulation, BF_10_ = 0.155, δ = −0.04.

Second, as an interaction was identified for a Cultural Group difference on the site-specific stimulation effect in general (not agent specific), we collapsed across agent and performed separate analyses for the dmPFC and rTPJ groups on total memory recognition scores. In the dmPFC group, a Stimulation Type x Cultural group interaction was identified, BF_10_ = 1.634, η_p_^2^ = 0.08. In the SEA group, anodal dmPFC stimulation reduced overall memory performance, BF_10_ = 4.824, δ = 0.68, whereas there was no overall change in the CA group, BF_10_ = 0.208, δ = −0.02.

Therefore, baseline performance on the SRE memory task was comparable between the groups and the subsequent effects of dmPFC stimulation in removing the SRE were comparable. However, an additional effect was identified in the SEA group, whereby dmPFC stimulation reduced overall memory in addition to removing the SRE. Stimulation of the rTPJ had no effects on SRE, or overall memory, in either group.

**Figure 3.**
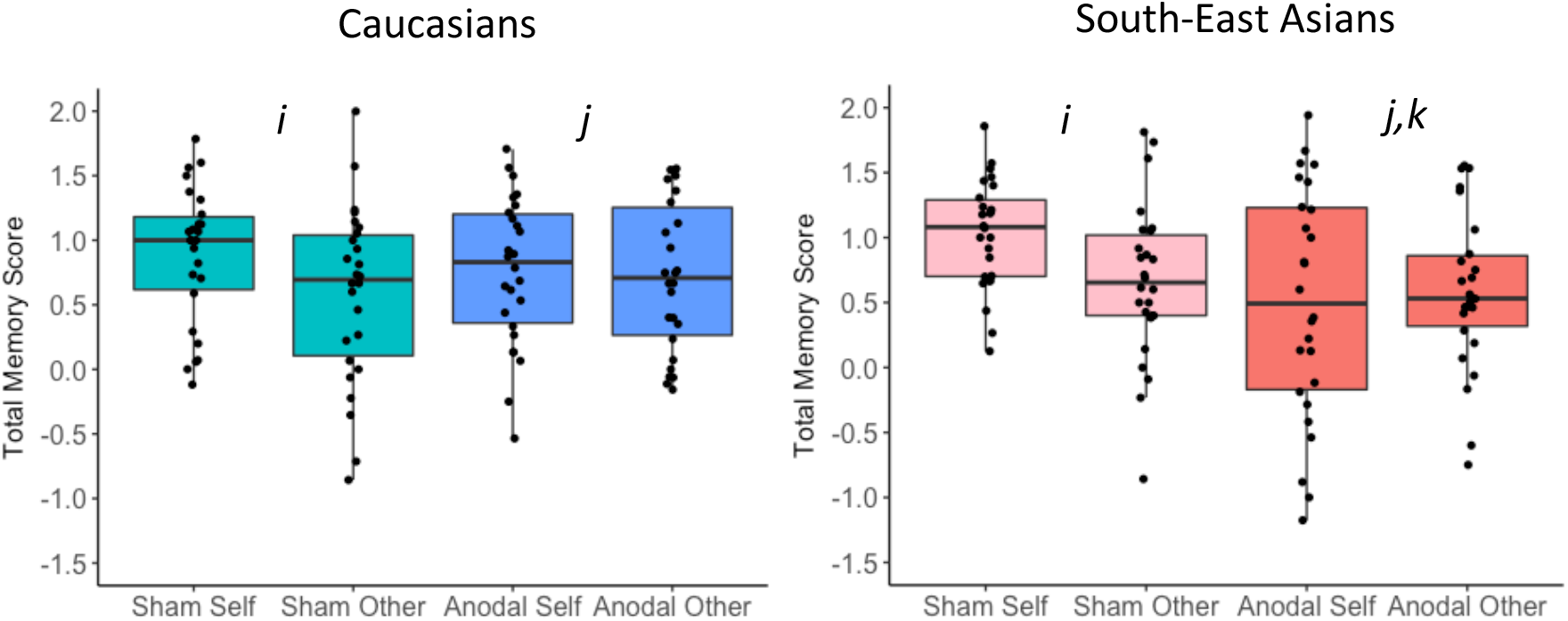
Self-reference effect (SRE) in episodic memory. *i)* A SRE was identified at baseline (sham-tDCS) with total memory score higher for self-encoded than other-encoded words in both groups (*i*), BF_10_ = 6.721; *ii)* After anodal stimulation to the dmPFC, the SRE was removed in both groups, BF_10_ = 6.542; *iii)* An additional reduction in overall (self & other) total memory score was identified in the SEA group only (*k*), BF_10_ = 4.824. The boxplot displays the median and the IQR. The whiskers extend to the most extreme datapoint within ±1.5*IQR.

### 3.3 Self-Reference Effect on Source Memory

No difference was identified between self and other encoding on source memories in the combined sample, BF_10_ = 0.185, η_p_^2^ = 0.004 and there was no interaction with cultural group, BF_10_ = 0.338, η_p_^2^ = 0.01. All stimulation main effect and interactions were in favour of the null (BF_10_ between 0.128-0.485).

Therefore, stimulation effects were only observed during the memory task with specific demands on self-referential encoding and not for the source memory component that did not explicitly rely on self-other encoding.

### 3.4 Association with Self-Construal Style

The exploratory analysis to identify whether stimulation effects differed by self-construal style in the SEA participants identified moderate evidence for a relationship between anodal HD-tDCS to the dmPFC induced reduction of the SRE and two dimensions of the self-construal scale; Self-Containment vs Connection to Others, r= 0.489, BF_10_= 9.678 and Self-direction vs Receptiveness to influence, r=0.457, BF_10_=6.763. Those who reported a greater interdependent self-construal style on these two dimensions, had greater reduction of the SRE after HD-tDCS to the dmPFC (see Figure 4). Evidence for the null model was identified for all stimulation effects on VPT measures (BF_10_ between 0.198-0.470).

**Figure 4.**
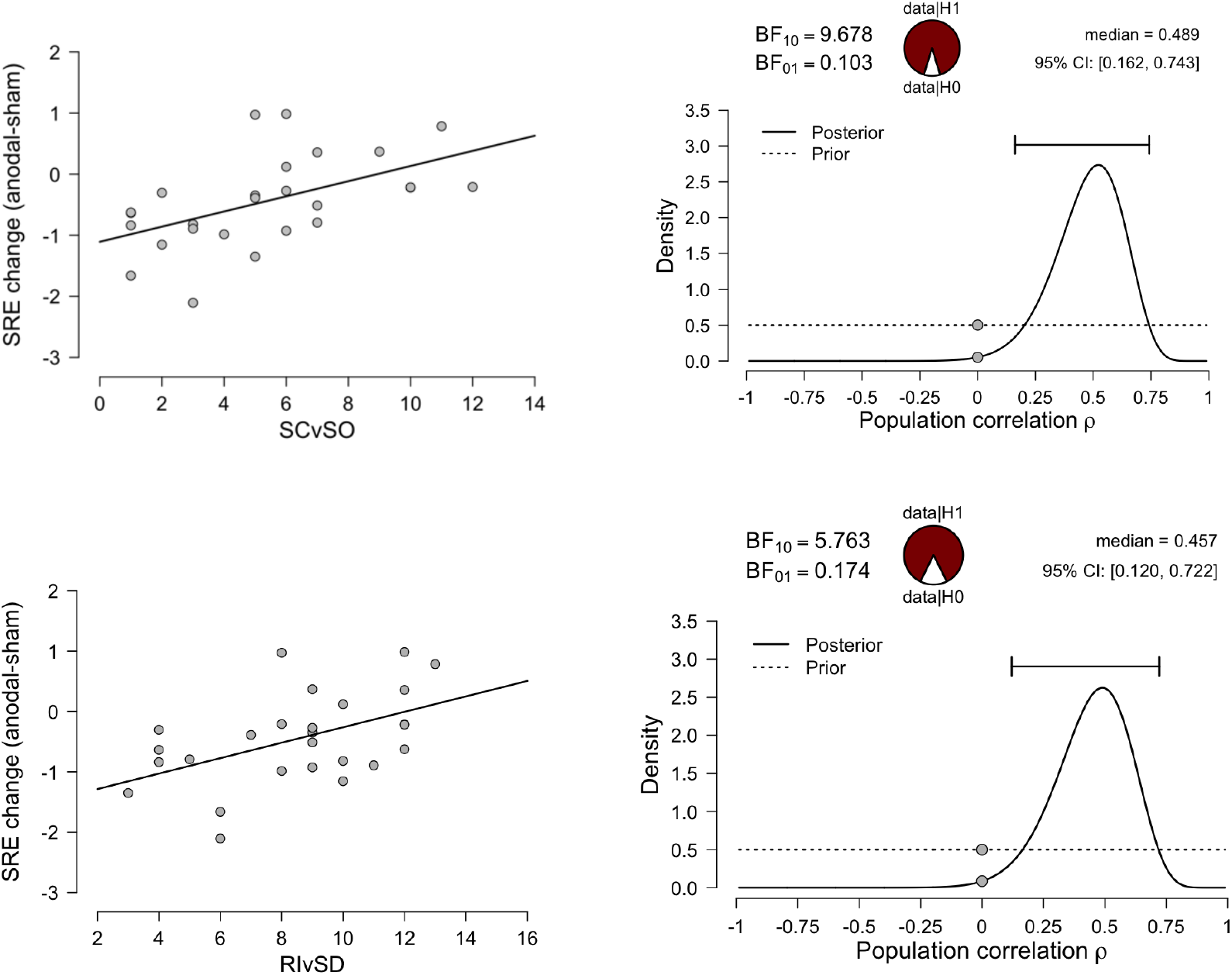
The effect of anodal stimulation of the dmPFC on the SRE correlated with self-construal style in the South-East Asian group. On both scales, lower scores represent a greater interdependent self-construal. Those who reported greater interdependence in the dimensions of Self-Containment vs Connection to Others (SOvSC) and Receptiveness to Influence vs Self-Direction (RIvSD) had the greatest reduction in SRE after anodal stimulation of the dmPFC. Prior and posterior distributions, the median effect size and a 95% credible interval are provided. The pie charts provide a visual representation of the evidence for the null or alternate model.

### 3.5 Adverse Effects, Mood Change, and Blinding

Adverse effects and mood change are detailed in Table 2. Stimulation type had no effect on positive mood change, BF_10_= 0.308, and there was no interaction with stimulation site, BF_10_= 0.228, and there was no further interaction with cultural group, BF_10_= 0.292. Likewise, there was no effect on negative mood change, BF_10_= 0.354, no interaction with stimulation site, BF_10_= 0.208, and no further interaction with cultural group, BF_10_= 0.615.

**Table 2.** Baseline cognition, depression, anxiety, and ASQ scores between Caucasian and East Asian subjects

|  | S-E Asians |  | Caucasian Australians |  | *Group*<br>*BF<sub>10</sub>* | *Site*<br>*BF<sub>10</sub>* | *Int*<br>*BF<sub>10</sub>* |
| --- | --- | --- | --- | --- | --- | --- | --- |
|  | dmPFC<br>(N=26) | rTPJ<br>(N=26) | dmPFC<br>(N=26) | rTPJ<br>(N=26) |  |  |  |
| International Shopping List | 29.0 (2.1) | 28.7 (3.0) | 29.2 (3.3) | 28.9 (3.1) | 0.22 | 0.24 | 0.27 |
| Identification Task | 2.70 (0.06) | 2.72 (0.07) | 2.71 (0.07) | 2.68 (0.06) | 0.28 | 0.23 | 1.79 |
| One-back Imn | 2.87 (0.09) | 2.88 (0.11) | 2.85 (0.08) | 2.86 (0.09) | 0.65 | 0.24 | 0.28 |
| Two-back Imn | 2.97 (0.11) | 2.99 (0.11) | 2.95 (0.09) | 2.93 (0.09) | 1.52 | 0.20 | 0.39 |
| Set-switching errors | 13.9 (5.8) | 17.0 (5.2) | 14.3 (5.3) | 19.0 (12.4) | 0.26 | 3.92 | 0.30 |
| CPAL errors | 30.0 (26.4) | 41.9 (35.2) | 42.4 (39.3) | 46.1 (32.9) | 0.42 | 0.39 | 0.30 |
| Socio-emotional cognition | 1.13 (0.09) | 1.14 (0.11) | 1.18 (0.11) | 1.13 (0.14) | 0.31 | 0.30 | 0.50 |
| ISL – delayed | 10.0 (1.3) | 10.3 (1.7) | 10.3 (1.4) | 10.2 (1.1) | 0.23 | 0.21 | 0.31 |
| Phonemic fluency | 14.9 (3.9) | 15.8 (4.5) | 17.0 (4.8) | 17.7 (5.7) | 1.58 | 0.29 | 0.27 |
| Semantic fluency | 23.5 (4.1) | 24.2 (3.9) | 25.5 (5.5) | 25.0 (5.8) | 0.57 | 0.21 | 0.33 |
| Stroop Effect | 22.3 (8.5) | 26.9 (11.8) | 22.0 (6.9) | 19.8 (8.6) | 1.31 | 0.24 | 1.16 |
| HADS depression | 3.3 (2.9) | 3.6 (2.3) | 2.3 (1.9) | 3.7 (2.8) | 0.33 | 0.67 | 0.46 |
| HADS anxiety | 6.7 (3.2) | 6.9 (3.6) | 7.2 (4.3) | 7.8 (4.5) | 0.29 | 0.23 | 0.30 |
| ASQ | 16.9 (5.7) | 18.9 (5.4) | 14.1 (5.4) | 14.6 (5.8) | 19.05 | 0.35 | 0.34 |

Imn= Log10 milliseconds speed of reaction for correct responses; CPAL= Continuous Paired Associates Learning; ISL= International Shopping List; NART= National Adult Reading Test; HADS= Hospital Anxiety and Depression Scale; ASQ= Autism Spectrum Quotient

Anodal stimulation resulted in more pronounced, albeit mild, adverse effects compared with sham, BF_10_= 3.195. However, this did not differ by stimulation site, BF_10_= 0.266 nor did it differ by cultural group, BF_10_= 0.330. There were no interactions between Stimulation Type and Stimulation Site, BF_10_= 0.266, Stimulation Type by Cultural Group, BF_10_= 0.219, nor Stimulation Type x Stimulation Site x Cultural Group, BF_10_= 0.528.

Participants were unable to predict which session was the active condition (59/104 correct guesses), BF_10_= 0.247. This did not differ by cultural group (30/52 & 29/52 correct guesses), BF_10_= 0.243.

### 3.6 Baseline Testing

Baseline cognitive performance is presented in Table 3. Cognitive differences were identified between the SEA and CA groups in two-back working memory, BF_10_= 1.52, phonemic fluency, BF_10_= 1.58, and Stroop Effect, BF_10_= 1.31. The SEA group also scored higher on the ASQ, BF_10_= 19.05. Those in the rTPJ study performed worse on a set-switching test, BF_10_= 3.92. However, none of these influenced the results, were dropped from the analyses, and are not discussed further.

**Table 3.**
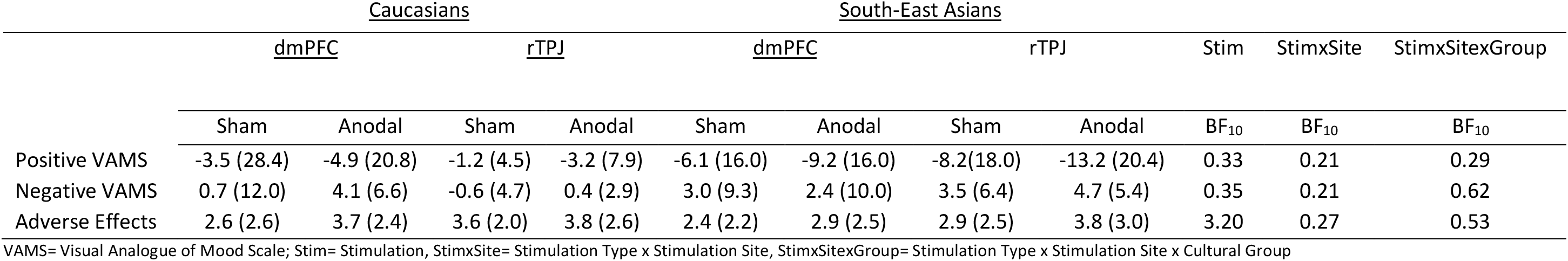
Summary of positive and negative mood ratings assessed pre and post stimulation for both Caucasian and East Asian cohorts. Mean difference between pre and post and standard deviations are reported

## 4. DISCUSSION

In the present study, we replicated key findings from a previous study (Martin, Huang, Hunold, & Meinzer, 2018) showing dissociable effects of HD-tDCS to the social brain. Specifically, a causal role for the dmPFC in self-referential memory and a causal role for the rTPJ in embodied mental rotation. Moreover, we provide the first evidence that HD-tDCS to the dmPFC has different effects on cognitive performance dependent on cultural background. As cultural background shapes the neural substrates supporting social cognition, the results provide further evidence that baseline differences in cognition are important considerations for brain stimulation effects.

One of the main findings of the present study was that, generally, when baseline performance between the two cultural groups was comparable, stimulation effects were subsequently comparable. On the other hand, when baseline performance differed, stimulation effects differed. This provides evidence for replicable effects of HD-tDCS on social brain functioning and a further consideration for minimising heterogeneity of tDCS response in future studies. Moreover, it provides further evidence for the notion that baseline cognitive functioning is an important consideration for subsequent stimulation effects (Benwell, Learmonth, Miniussi, Harvey, & Thut, 2015; Looi et al., 2016; Martin, Meinzer, et al., 2017; McConathey et al., 2017; Sarkar, Dowker, & Cohen Kadosh, 2014; Turkeltaub et al., 2012). Both cultural groups were comparable at baseline when taking the allocentric perspective during the level two VPT task and anodal stimulation to the rTPJ reduced the egocentric interference in both groups. Likewise, both groups had a comparable SRE for episodic memory and anodal stimulation to the dmPFC removed this bias. Unique effects were identified at baseline for the level one VPT task, with the SEA group influenced by the alternate perspective to a greater extent than the CA group. Stimulation of the dmPFC shifted the performance of CA closer to that of the SEA during the egocentric perspective trials by increasing the interference from the allocentric judgement. However, stimulation had no effect on the performance of SEA. Anodal stimulation to the dmPFC also had a unique effect in the SEA cohort of reducing overall memory performance, whereas in the CA, overall memory performance remained the same and only the bias towards self-referential memories was removed.Previous studies have highlighted differential tDCS effects on the social brain based on sex (Adenzato et al., 2017; Martin, Huang, et al., 2017), but this is the first study to provide evidence that cultural background may also be a necessary consideration. Although this effect may be limited to social cognition where culture is a recognised source of variance, performance on other cognitive domains has been shown to be culturally specific (Nisbett & Miyamoto, 2005) and should be considered in future brain stimulation studies.

The results of the current study provide evidence for replicable and dissociable effects of HD-tDCS to regions within the social brain on self-other processing and provides further evidence that baseline performance is an important consideration for subsequent tDCS effects. With the current focus on improving the replication of results in psychology (Open Science, 2015), the replicable effects of social brain stimulation provide strong evidence for regional and task specific roles of the dmPFC and rTPJ in self other processing. Also, replicated robust effects of HD-tDCS helps dispel claims regarding the effectiveness of tDCS to have any meaningful effect on cognition (Horvath, Forte, & Carter, 2015). Replicating effects in a separate cohort from a different cultural background goes some way to address issues concerning homogeneity of conclusions based on limited participant sampling from predominantly Western backgrounds (Henrich, Heine, & Norenzayan, 2010). Moreover, the field of cultural neuroscience has emerged out of a greater need to incorporate culture into our understanding of how brain development is shaped by a complex and dynamic interplay between biology and socialisation through different societal lenses (Park & Huang, 2010). The current study, to the authors knowledge, is the first to extend this framework to the field of brain stimulation presenting a promising avenue for future cultural neuroscientific research.

Previously it has been shown that people from East Asian cultural backgrounds are more integrative in their approach to level one VPT tasks, as indexed by a greater influence of alternate perspectives during egocentric perspective taking (Kessler et al., 2014; Wu & Keysar, 2007). We provide further support for this cultural difference in the current study, although we extend this evidence to show that the alternate perspective (egocentric or allocentric) interfered to a greater extent in SEA compared with CA and not solely during the egocentric perspective task as previously demonstrated. Our findings are consistent with previous research showing that East Asians are more holistic in their cognitive style, whereby the context exerts more influence on the current task (Nisbett, Peng, Choi, & Norenzayan, 2001). In the present study, a holistic style would manifest as greater salience or awareness, and subsequent interference, of the alternate perspective, as demonstrated in the SEA participants. As anodal stimulation of the dmPFC increased the influence of allocentric information on egocentric judgements in CA, performance was shifted closer to that shown at baseline by the SEA group during egocentric perspectives. The lack of stimulation effects in the SEA group suggests that either the dmPFC is more active in the SEA such that stimulation has no differential physiological effect or that underlying brain-behaviour associations are different between the two groups. The fact that stimulation shifts performance in the CA towards the SEA provides evidence for the former, which suggests that the CA recruit the dmPFC to a lesser extent during these types of tasks. As the dmPFC is consistently associated with tasks requiring integration of social information (Brosch, Schiller, Mojdehbakhsh, Uleman, & Phelps, 2013; Ferrari et al., 2016; Wittmann et al., 2016), CA may adopt strategies that are less reliant on integration of information and therefore less reliant on dmPFC recruitment, consistent with previous research showing that Westerners are less relational and more rigid in their sense of self regardless of the situational demands (Varnum, Grossmann, Kitayama, & Nisbett, 2010). Further research, especially incorporating concurrent neuroimaging and functional MRI, may further elucidate causal brain-behaviour relationships that differ in groups from diverse cultural backgrounds.

Although SRE was removed after anodal stimulation of the dmPFC, a replication of the effects seen in CA participants, anodal stimulation to the dmPFC had an additional effect of reducing overall memory in the SEA cohort. Although the mPFC shows a self-other gradient from ventral to dorsal regions, there is evidence that this is less demarcated in people from East Asian cultural backgrounds (Harada et al., 2010) coupled with less mPFC activity associated with self-referential processing (Ma, Bang, et al., 2014). As dmPFC is associated with a wide range of memory processes (Euston, Gruber, & McNaughton, 2012), differences in underlying neural brain-behaviour association may result in the differences in stimulation response. As functional connectivity patterns with the dmPFC are also culturally specific (Li et al., 2018), connectivity differences between the cultural groups may also be relevant for the culturally specific results and should be considered in future research using concurrent tDCS and fMRI (Gbadeyan et al., 2016).

The exploratory analyses correlating stimulation response with self-construal provide a cautionary note that culture is not a binary construct. Independent and interdependent self-construal exists on a continuum and variation exists both between and within cultures (Vignoles et al., 2016). Although not assessed in the present study, similar variation will exist in Western cohorts and requires examination in future studies. Heterogeneity in stimulation response provides both an opportunity for the scientific method of explaining individual differences and a challenge for clinical applications that aspire to show consistent effects of brain stimulation that are widely applicable. However, further knowledge concerning mediating factors that may improve stimulation response will ultimately improve the application in both experimental and clinical contexts. This is of particular relevance for future studies in clinical cohorts with social cognitive impairments such as psychosis and autism. Future research is necessary to extend this research to other cultural groups. It also should be explored if culture mediates stimulation response in other cognitive domains and at other stimulation sites. Likewise, concurrent tDCS-fMRI will improve our understanding of the basic neurophysiological effects of HD-tDCS, how it affects underlying neural tissue, and how cultural background may modulate stimulation effects.

In sum, this is the first study to demonstrate replicable effects of HD-tDCS to the social brain as well as unique differences in social brain stimulation response in people from a different cultural background. The replicated effects provide robust evidence that HD-tDCS can consistently influence social cognition. The culturally specific effects provide evidence that HD-tDCS is a useful method for investigating cultural differences in the underlying social brain and its relationship to self-construal.

## Contributions

AM and MM conceived of the study. AM designed the study. Testing and data collection were performed by AM and PS. AM performed the data analysis. AM drafted the manuscript and MM and PS provided critical revisions. All authors approved the final manuscript for submission.

## Acknowledgements

We would like to acknowledge Dr Ilvana Dzafic for assistance with coding the tasks and Jasmine Huang for data collection. The study was supported through a Future Fellowship [FT120100608] and a strategic seed-funding grant from the University of Queensland, awarded to Marcus Meinzer.

## Notes

#### Summary of Updates

Revised

